# Investigating how *Salmonella* colonise alfalfa using a whole genome screen

**DOI:** 10.1101/2023.10.24.563821

**Authors:** Emma R. Holden, Haider Al-Khanaq, Noémie Vimont, Mark A. Webber, Eleftheria Trampari

**Affiliations:** Quadram Institute Bioscience, Norwich Research Park, Norwich, Norfolk, NR4 7UQ, U.K.; Norwich Medical School, University of East Anglia, Norwich Research Park, Norwich, Norfolk, NR4 7TJ, U.K.

**Author notes:** Corresponding author: Eleftheria Trampari.

**Keywords:** TraDIS, foodborne pathogens, fresh produce, Salmonella, functional genomics, food safety

## Abstract

Enteropathogenic bacteria including *Salmonella* regularly cause outbreaks of infection from fresh produce posing a significant public health threat. *Salmonella*’s ability to persist on fresh produce for extended periods is partly attributed to its capacity to form biofilms, which poses a challenge to food decontamination and facilitates persistence in the food chain. Preventing biofilm formation on food products and in food processing environments is crucial for reducing the incidence of foodborne diseases. Understanding the mechanisms of colonisation and establishment on fresh produce will inform the development of decontamination approaches. We used Transposon-directed Insertion site sequencing (TraDIS-*Xpress*) to investigate the mechanisms employed by *Salmonella* enterica serovar Typhimurium to colonise and establish itself on fresh produce at critical timepoints following infection. We established an alfalfa infection model and compared the findings to those obtained from glass surfaces. Our research revealed dynamic changes in the pathways associated with biofilm formation over time, with distinct plant-specific and glass-specific mechanisms for biofilm formation, alongside the identification of shared genes playing pivotal roles in both contexts. Notably, we observed variations in the significance of factors such as flagella biosynthesis, lipopolysaccharide (LPS) production, and stringent response regulation in biofilm development on plant versus glass surfaces. Understanding the genetic underpinnings of biofilm formation on both biotic and abiotic surfaces offers valuable insights that can inform the development of targeted antibacterial therapeutics, ultimately enhancing food safety throughout the food processing chain.

**Funding:** The authors gratefully acknowledge the support of the Biotechnology and Biological Sciences Research Council (BBSRC); ERH, JAA, HAK, MAW and ET were supported by the BBSRC Institute Strategic Programme Microbes and Food Safety BB/X011011/1 and its constituent project BBS/E/F/000PR13635. NV was supported by the Food Safety Research Network grant BB/X002985/1 awarded to ET.

**Data availability:** Nucleotide sequence data supporting the analysis in this study has been deposited in ArrayExpress under the accession number E-MTAB-13495. The authors confirm all supporting data, code and protocols have been provided within the article or through supplementary data files.

## Introduction

Enteropathogenic bacteria present an evolving threat to public health. Historically, these pathogens were predominantly linked to meat products. However, in recent years, fresh produce is emerging as a major source of these outbreaks, being implicated in over a third of reported outbreaks in certain countries (Brennan et al., 2022). The majority of these outbreaks are associated with ready-to-eat crops, although some cases have been attributed to the mishandling of vegetables that are typically subjected to cooking processes (Launders et al., 2016). Certain human pathogens, such as *Salmonella*, are able to colonise various ecological niches and survive outside their primary host (Humphrey, 2004). *S. enterica* has been implicated in numerous recent multistate outbreaks in the USA associated with contaminated fruits and vegetables, including lettuce, tomatoes, alfalfa, cucumbers, and melons (Heaton and Jones, 2008, EFSA et al., 2017). Recent studies have demonstrated *Salmonella*’s ability to actively colonise plant tissues employing specific mechanisms (Salazar et al., 2013) and *Salmonella* has been found to persist in produce for extended periods, with viability lasting over six months (Islam et al., 2004).

*Salmonella*’s adaptive strategy to persist in the challenging plant environment includes the formation of biofilms. Biofilms are structured, aggregated communities of microorganisms encased in an extracellular matrix and attached to surfaces (Monier and Lindow, 2005). These communities play a critical role in enabling pathogenic bacteria to adhere to fresh produce increasing the risk of enteric disease transmission (Yaron and Römling, 2014). Bacteria within biofilms exhibit intrinsic tolerance to high concentrations of antimicrobials, biocides, and disinfectants, which complicates decontamination efforts and poses challenges for ensuring food safety. Previous studies have contributed valuable insights into the mechanisms underlying *Salmonella’s* biofilm formation and its ability to persist on plants, highlighting the significance of these processes in the context of food safety and public health (Fett, 2000, Brandl, 2006, Brandl and Mandrell, 2002). However, the range of plants and conditions studied has been limited and little whole genome analysis of factors responsible for plant colonisation has been done to date.

Transposon sequencing (TnSeq) approaches have previously been used to determine the mechanisms through which bacteria survive in different environments. For example, TnSeq was used to identify the genes involved in *Pseudomonas simiae* colonisation of plant roots, which highlighted the importance of genes involved in flagella production, cell envelope biosynthesis, carbohydrate metabolism and amino acid transport and metabolism (Cole et al., 2017). We have previously used another TnSeq variant, TraDIS-*Xpress*, to identify genes involved in biofilm formation in *Escherichia coli* (Holden et al., 2021) and *Salmonella enterica* serovar Typhimurium (Holden et al., 2022) on glass over time. TraDIS-*Xpress* builds on conventional transposon sequencing approaches by using larger denser transposon mutant libraries and by incorporating an outwards-transcribing promoter into the transposon element (Yasir et al., 2020). Induction of this promoter enables increased expression of genes downstream of transposon insertions thereby facilitating investigation into how expression, as well as gene disruption, affects survival of the mutant in a given condition. This approach also generates information about essential genes which do not tolerate insertional inactivation by transposons and can therefore not be assayed with conventional tools.

In this study, we established an alfalfa plant infection model that was used in conjunction with TraDIS-*Xpress* to investigate gene essentiality in biofilm formation on alfalfa over time. A comprehensive library of *S*. Typhimurium transposon mutants was cultivated on sprouted alfalfa plants and mutants were isolated at different stages to identify the genes involved in biofilm development *in planta* over time. Comparisons were made with data from biofilms on glass surfaces (Holden et al., 2022). This allowed for the identification of plant-specific and glass-specific mechanisms used by *S*. Typhimurium to establish in organic and inorganic surfaces, as well as conserved genes that play crucial roles on both surfaces.

We showed different sets of genes were needed for colonisation of alfalfa compared to glass, with variations in the importance of factors including flagella biosynthesis, LPS production, and stringent response regulation in biofilm development on plants. Understanding the genes involved in colonisation and biofilm formation on both biotic and abiotic surfaces will provide valuable insights for the development of targeted antibacterial therapeutics to enhance food safety throughout the food processing chain.

## Results

### Establishment of an alfalfa plant infection model

To assess *S. Typhimurium*’s ability to establish and proliferate on plant hosts, an alfalfa seedling infection model was established (Figure 1). Alfalfa was chosen as an important vehicle of *Salmonella* infection and as an easily studied laboratory model. Initially, seeds underwent sterilisation and were allowed to germinate in Murashige-Skoog (MS) medium for three days (Figure 1-A). Following this germination period, the seedlings were inoculated at the root-shoot intersection with a *S. Typhimurium* strain marked either with the *lacZ* marker (*14028S::lacZ*) for blue colony selection and counting; or with a *lux* tagged strain for live visualisation on seedlings (*14028S::lux*) (see Figure 1D).

**Figure 1.**
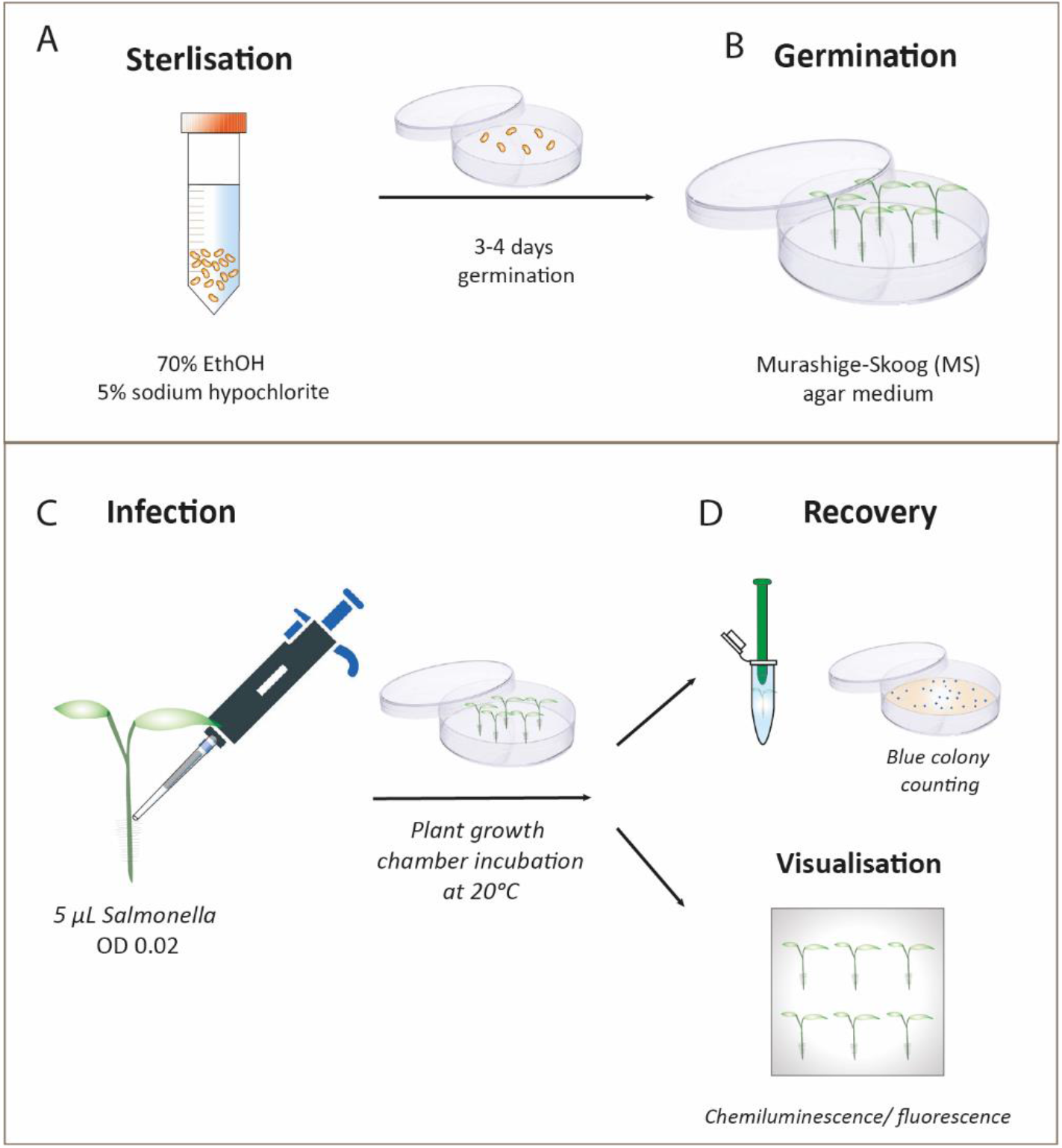
Alfalfa Plant Infection Model. A. Alfalfa seeds were sterilised by immersion in 70% ethanol for 30 seconds, followed by a 3-minute wash in 5% sodium hypochlorite. B. Subsequently, the sterilised seeds were left to germinate in darkness at 20°C in Murashige-Skoog (MS) agar medium for 3-4 days. C. Infection of the seedlings was performed at the root-shoot intersection using 5 μL of Salmonella inoculum, normalised to an optical density (OD) of 0.02. Infected seedlings were then transferred to fresh MS plates and incubated in a benchtop plant growth chamber at 20°C. D. To facilitate selection via blue colony screening, Salmonella recovery and quantification were performed using the 14028S::lacZ strain after 6, 24, and 48 hours. Inoculated seedlings were homogenised by mechanical disruption using a pestle to release the bacterial cells. Cell suspensions were subjected to serial dilution and plated onto X-gal/IPTG LB plates for further analysis. Visualisation experiments were conducted using the 14028S::lux strain to monitor the localisation of Salmonella on the seedlings via chemiluminescence. For microscopy, the 14028S:mplum strain was

### *Salmonella* effectively colonises alfalfa sprouts and increases in numbers over time

To investigate the effectiveness of *Salmonella* colonisation in alfalfa seedlings, a chemiluminescence-tagged strain of *Salmonella* (*14028S::lux*) was employed to infect seedlings three days after germination. Following infection, the seedlings were washed in PBS and were subsequently visualised using a Gel documentation system (Biorad). Notably, specific colonisation of the roots by *Salmonella* was observed, even from very early stages post infection (3 hours) with an evident increase in presence over time, as indicated by the chemiluminescence intensity (Figure 2A). To quantify the *Salmonella* load on the seedlings over time, a strain tagged with *lacZ* (*14028S::lacZ*) (Holden et al., 2020) was utilised to facilitate selection and counting. Cells were recovered after 8, 24, 48 and 72-hours growth, demonstrating a significant increase in *S*. Typhimurium colonisation of alfalfa over time (see Figure 2B

**Figure 2.**
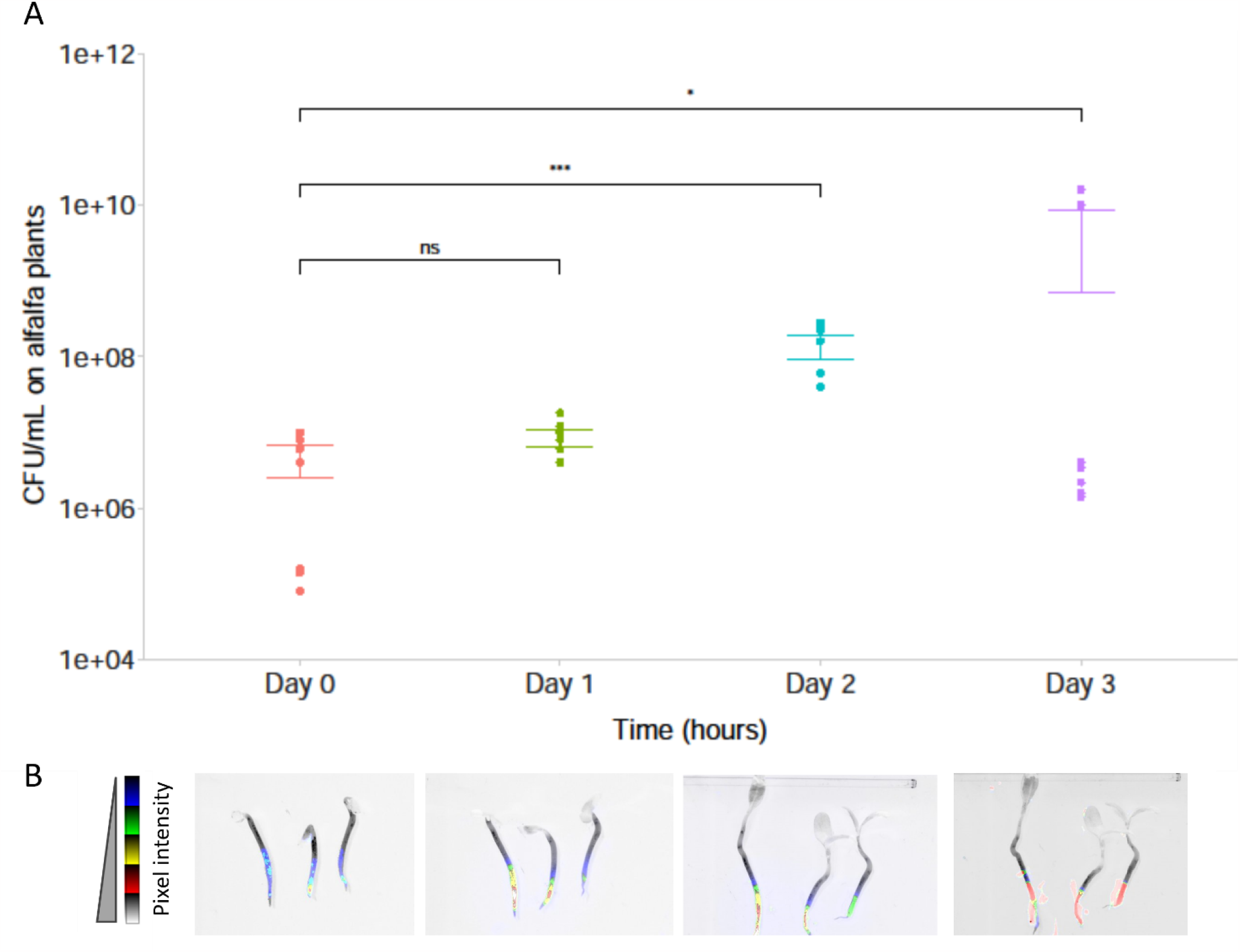
Salmonella effectively colonises the alfalfa model. A. Number of cells recovered (per seedling) of 14028S::lacZ followed by at 8 hours, 24 hours, and 48 hours post-infection demonstrating a significant increase in Salmonella numbers over time. Each spot represents data from an independent replicate seedling. B. Visualisation of 14028S::lux on alfalfa seedlings through chemiluminescence monitoring at 3 hours, 24 hours, and 48 hours post-infection, demonstrating the specific seedling colonisation by Salmonella over time. Chemiluminescence is depicted using five colors, with the transition from blue to red indicating higher intensity and hence cell density.

### Genes involved in *Salmonella* establishment on alfalfa over time

TraDIS-*Xpress* was used to identify genes involved in alfalfa colonisation by *S*. Typhimurium over 3 days (24-, 48- and 72-hours post-seeding). These timepoints were chosen to capture the potentially diverse mechanisms required by *Salmonella* at different stages of alfalfa colonisation. This includes the early stages involving initial attachment and microcolony formation (at 24 hours, representing Day 1) and the subsequent phases of establishment and biofilm formation (spanning 48 to 72 hours, representing Day 2 and 3). We identified 69 genes as significantly involved in *S*. Typhimurium colonisation and biofilm formation on alfalfa sprouts over time (supplementary table 1). These included genes involved in LPS biosynthesis, DNA housekeeping, respiration and responding to stress (figure 3). Variation in insertion frequency per gene between replicates was low, indicating minimal experimental error (supplementary figure 1).

**Figure 3.**
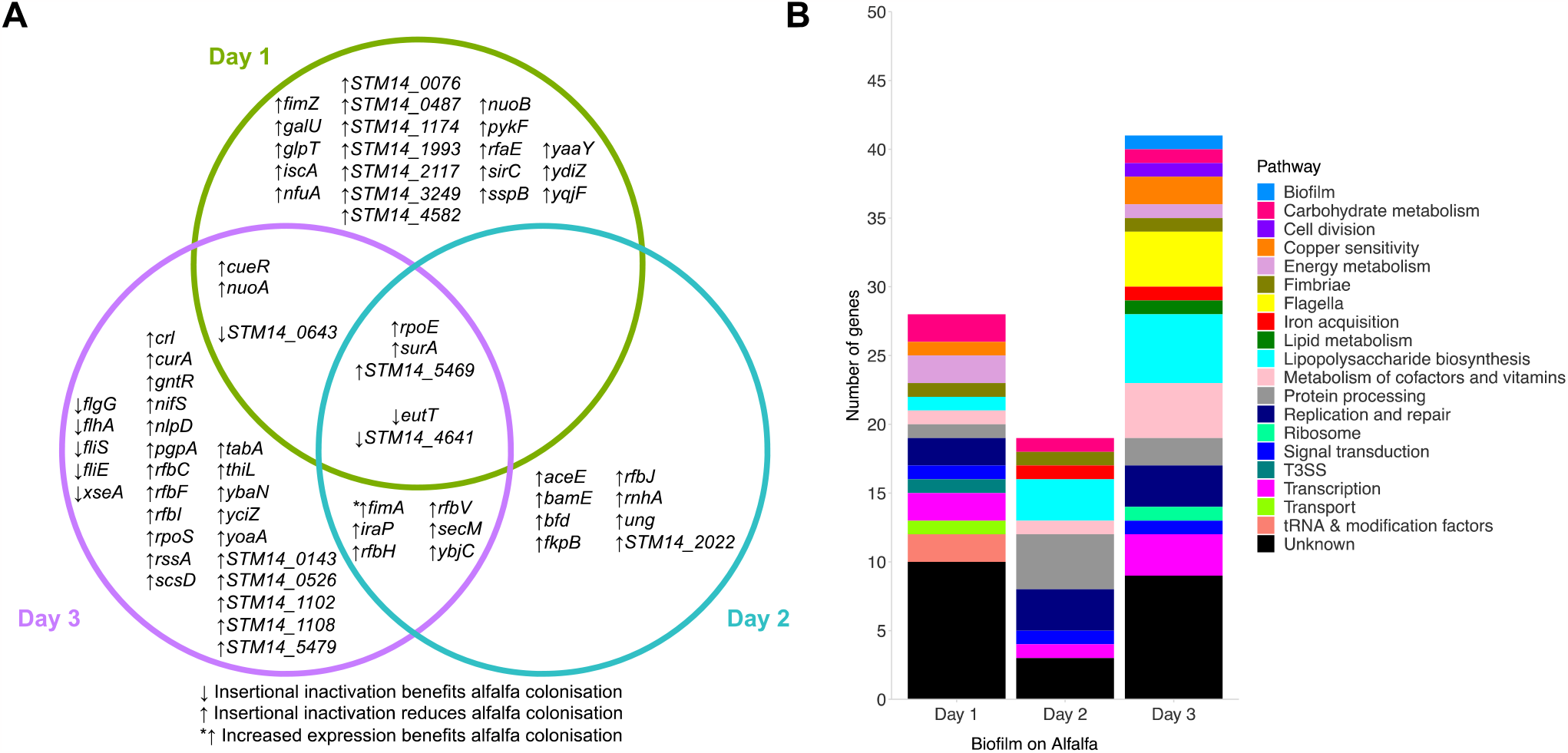
A) Genes and B) pathways identified by TraDIS-Xpress to be involved in alfalfa colonisation over time.

Genes involved in adhesion were identified as beneficial after 24 hours growth, including previously reported genes, such as a negative fimbrial regulator *fimZ* (Saini et al., 2009) and type III secretion system component *sirC* (Rakeman et al., 1999). After 48 hours, genes involved in DNA housekeeping (*rnhA* and *ung*) (Ogawa and Okazaki, 1984, Duncan et al., 1978),(Ogawa and Okazaki, 1984, Duncan et al., 1978), iron storage (*bfd*) (Quail et al., 1996) and outer membrane protein assembly (*bamE*) (Sklar et al., 2007) benefitted the further establishment of *Salmonella* on alfalfa. Following 72 hours growth, genes associated with mature biofilm formation were identified as being important, including those with roles in LPS O-antigen production (*rfbF, rfbI, rfbC, rfbV* and *rfbH*) (Wang et al., 2015), flagella biosynthesis (*flgG, flhA, fliS* and *fliE*) (Macnab, 1992) and responding to stress (*rpoS, iraP* and *crl*) (Battesti et al., 2011).

Five genes were shared among all the time points tested; these were *eutT, surA, rpoE, STM14_4641* and *STM14_5469*. Loss of function of the *eut* operon through disruption of *eutT* (Penrod and Roth, 2006) was predicted to be beneficial to *S*. Typhimurium establishment at all time points tested. Transcription of the RNA-directed DNA polymerase *STM14_4641* was detrimental to colonisation throughout its growth on alfalfa sprouts. There were fewer transposon mutants across all three days in *surA* (outer membrane protein chaperone (Lazar and Kolter, 1996)),(Lazar and Kolter, 1996)), *rpoE* (sigma factor involved in responding to misfolded protein stress (Alba and Gross, 2004))(Alba and Gross, 2004)) and *STM14_5469* (unknown function), which suggests these genes are beneficial throughout all stages of alfalfa colonisation.

### Conserved pathways crucial for biofilm formation on alfalfa sprouts and glass

We have previously identified genes essential for biofilm formation on glass over time using the same *S*. Typhimurium TraDIS library (Holden et al., 2022). Insertion frequencies in mutant libraries colonising glass or plant surfaces were both compared to planktonic cultures grown for the same amount of time. This acted as a standard to demonstrate where transposon insertions affected surface colonisation relative to planktonic growth, and the subsequent gene lists were then compared. This found some core pathways involved in *S*. Typhimurium establishment on both surfaces which included flagella biosynthesis, LPS production, respiration, iron storage and stress responses. Seven genes were found to be conserved between biofilms grown on alfalfa sprouts and on glass (figure 4). These were *nuoA* and *nuoB*, involved in synthesis of the first NADH hydrogenase in the electron transport chain (Archer and Elliott, 1995),(Archer and Elliott, 1995), fimbrial subunit *fimA* and its regulator *fimZ* (Saini et al., 2009), *rfbJ* involved in LPS O-antigen synthesis (Wang et al., 2015), *ybaN* predicted to have a role in iron acquisition (Seo et al., 2014), and stress response sigma factor *rpoS* (Gentry et al., 1993).(Gentry et al., 1993). The ethanolamine utilisation pathway played an important role in *S*. Typhimurium establishment on both alfalfa sprouts (*eutT*) and on glass (*eutQ*) at all time points tested, with disruption of each gene seen to aid colonisation. Together, this reveals a core set of pathways involved in colonisation of both biotic and abiotic surfaces.

**Figure 4.**
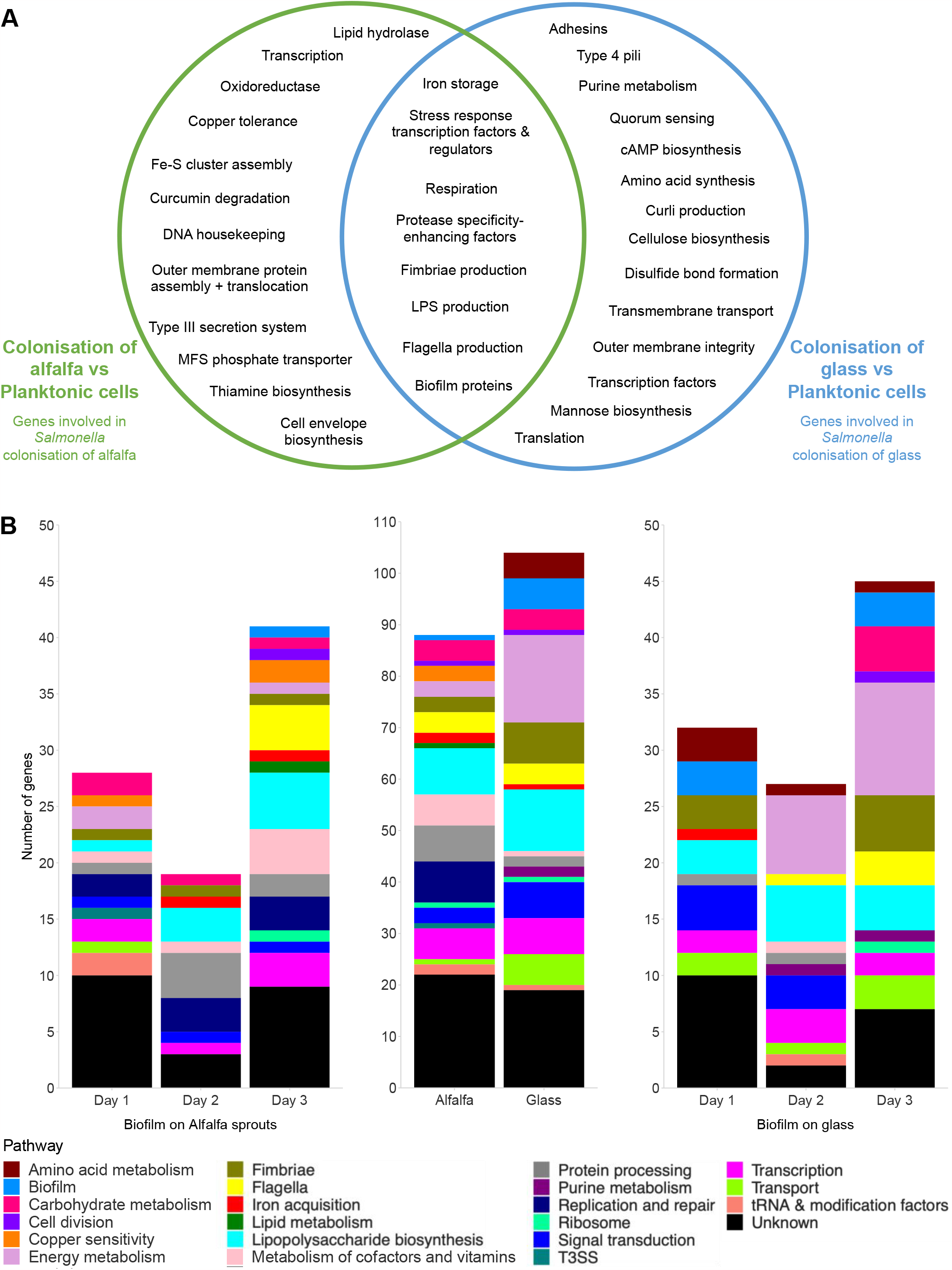
A) Conserved and surface-specific pathways involved in S. Typhimurium colonisation of alfalfa sprouts and glass. B) Abundance of genes in each pathway at over time for biofilms grown on alfalfa sprouts or glas

### Differential Flagella and Lipopolysaccharide Biosynthesis on Alfalfa vs. Glass

Deletion mutants were constructed in targets identified by TraDIS-*Xpress* to investigate their effects on colonisation and establishment on diverse surfaces (both inorganic and organic). These mutants were then subjected to competitive colonisation experiments with wild type *S*. Typhimurium strains on both glass and alfalfa surfaces. Equal numbers of mutant and wild type were inoculated onto glass beads and alfalfa plant sprouts. Subsequently, the percentage change in mutants within the recovered populations from each surface was determined over time.

TraDIS-*Xpress* indicated that inactivation of genes involved in flagella biosynthesis was beneficial for plant colonisation after 72 hours (Figure 5A). As flagella are detected by the plant’s immune system, aflagellated cells may have a competitive advantage in these communities once an immune response is raised. Our previous work found aflagellated cells were disadvantaged at colonising glass surfaces (Holden et al., 2022). To characterise the role of flagella in *S*. Typhimurium establishment on both environments, a deletion mutant of the main flagella biosynthetic regulator (*flhDC*) and a component of the flagella export machinery (*flhA*) were grown on glass and alfalfa sprouts in competition with wild type *S*. Typhimurium. At the initial stages of colonisation (Day 1), Δ*flhDC* and Δ*flhA* exhibited a significantly enhanced competitive advantage at colonising glass but were competitively disadvantaged at colonising alfalfa plants (Figure 5B). This contrasts with the prediction from the TraDIS-*Xpress* findings, however TraDIS measures competitive fitness in a large pool of competing mutants where individuals lacking flagella may benefit from adhesion of neighbouring cells whilst also being less susceptible to host immunity.

**Figure 5.**
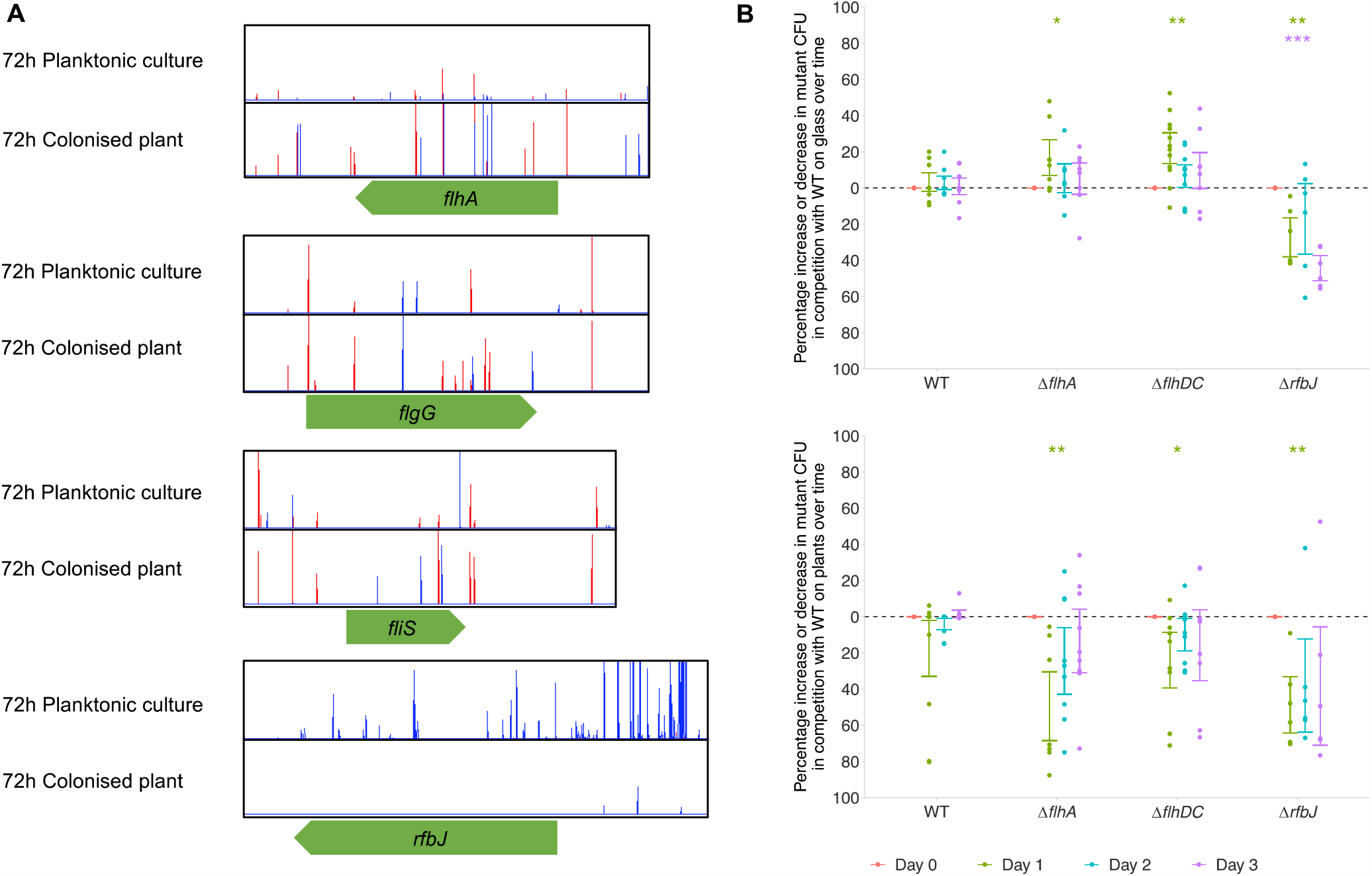
A) Insertion loci and frequency in and around genes involved in flagella biosynthesis (flhA, flgG and fliS) and LPS O-antigen biosynthesis (rfbJ) following growth on alfalfa sprouts relative to planktonic growth. Red lines indicate the transposon-located promoter is facing left-to-right and blue lines indicate it is oriented right-to-left. Images are representative of two independent replicates. B) Percentage increase or decrease in flhA, flhDC and rfbJ deletion mutants in biofilms formed on glass (top panel) and alfalfa plant sprouts (bottom panel) in competition with wild type (WT) S. Typhimurium. Points show changes in the percentage of mutant CFU relative to time point 0 and show 3 technical and 4 biological replicates. Error bars denote 95% confidence intervals and asterisks show significant differences (One-sample t-test, change from 0) of each mutant from time point 0, where time points are distinguished by colour: * p < 0.05, ** p < 0.01, *** p < 0.001, **** p < 0.0001.

LPS core and O-antigen biosynthesis genes were beneficial for growth on alfalfa sprouts, however the impact of different LPS biosynthesis genes on *S*. Typhimurium colonisation varied. Some exhibited beneficial effects when inactivated during glass colonisation, while others had detrimental impacts. Based on the TraDIS-*Xpress* data, *rfbJ* was beneficial for growth and establishment on alfalfa sprouts, whereas inactivation of the gene was beneficial for establishment on glass. We created a deletion mutant of *rfbJ* in *S*. Typhimurium to investigate its effect on glass and plant colonisation. Deletion of *rfbJ* resulted in reduced colonisation of both glass and plant over time (Figure 5B). This confirmed the predicted importance of this gene for adhesion and colonisation of both surfaces.

### Genes involved in copper tolerance, type III secretion regulation and curcumin degradation conferred a competitive advantage to *Salmonella* establishment on alfalfa

Analysis of the TraDIS-*Xpress* data found pathways involved in *S*. Typhimurium establishment on alfalfa plants that were not involved during biofilm formation on glass. These included type III secretion regulation (*sirC*) (Rakeman et al., 1999) and Fe-S cluster assembly (*iscA*) (Vinella et al., 2009),(Vinella et al., 2009), which were beneficial at the early stages of colonisation of alfalfa, curcumin degradation (*curA*) (Hassaninasab et al., 2011) was beneficial following 72 hours growth on alfalfa and copper tolerance (*cueR*) (Osman et al., 2010) was beneficial following 24 and 72 hours growth on alfalfa.

Gene deletion mutants were made in these genes and grown in the presence of wild type *S*. Typhimurium on glass and alfalfa plants to investigate their effects on colonisation. Deletion of *iscA* resulted in a competitive disadvantage for colonisation of both glass and alfalfa plants, supporting the TraDIS-*Xpress* findings (figure 6B). Deletion of *cueR* caused a competitive disadvantage in colonisation of alfalfa plants, but there was no significant change in glass colonisation, demonstrating that expression of *cueR* is only beneficial for colonisation of plant surfaces and not glass surfaces. There was no significant change in the percentage of Δ*sirC* or Δ*curA* mutants in the biofilm over time on either glass or plants, suggesting the effects of these genes on colonisation observed in the TraDIS-*Xpress* data cannot be quantified by this assay.

**Figure 6.**
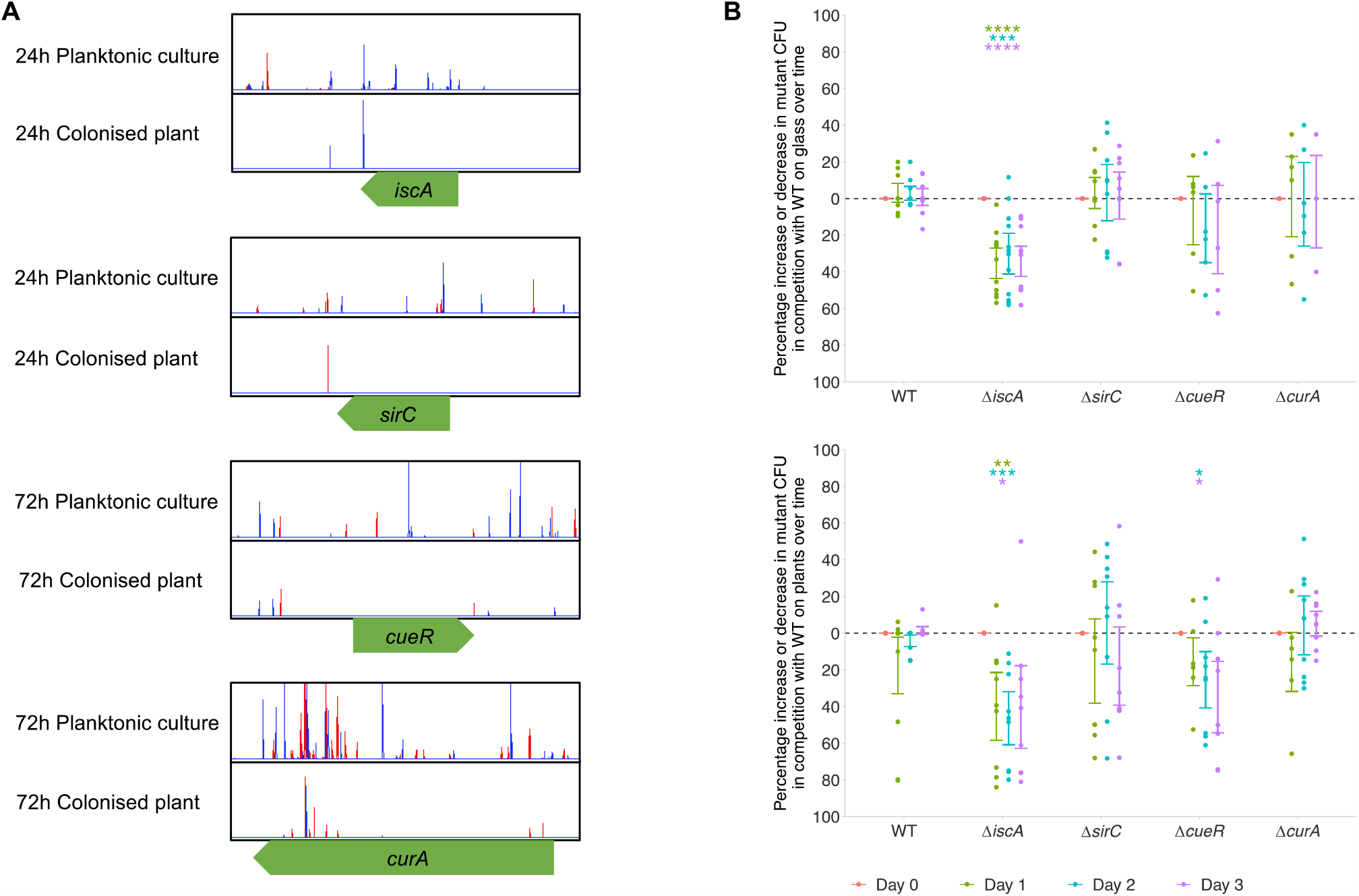
a) Transposon insertions within and around iscA, sirC, cueR and curA in S. Typhimurium planktonic culture compared to biofilms on an alfalfa plant grown for 24 or 72 hours. Lines show the insertion loci and the height of the lines shows the number of reads mapped to the loci. The colour of the line indicates the orientation of the promoter within the transposon: red lines denote the promoter is promoting transcription left-to-right, and blue lines denote right-to-left. Plot files shown are representative of two independent replicates. b) Percentage increase or decrease in iscA, sirC, cueR deletion mutants in biofilms formed on glass (top panel) and alfalfa plant sprouts (bottom panel) in competition with wild type (WT) S. Typhimurium. Points show changes in the percentage of mutant CFU relative to time point 0 and show 3 technical and 4 biological replicates. Error bars denote 95% confidence intervals and asterisks show significant differences (One-sample t-test, change from 0) of each mutant from time point 0, where time points are distinguished by colour: * p < 0.05, ** p < 0.01, *** p < 0.001, **** p < 0.0001.

## Discussion

The primary objective of this study was to identify the mechanisms employed by *S*. Typhimurium to colonise effectively and establish on fresh produce and compare these to the ones required for colonisation and biofilm formation on glass, across various stages of colonisation. To achieve that, we established a fresh produce alfalfa infection model and draw a comparative analysis using genome-wide transposon sequencing (TraDIS-*Xpress*) on alfalfa, comparing them to mechanisms previously identified for biofilm formation on glass surfaces (Holden et al., 2022). Our aim was to discern the extent to which these mechanisms are universally necessary for adhesion, colonisation, and establishment on organic surfaces in contrast to inorganic ones. Our working hypothesis was that there would be both common and bespoke mechanisms important for colonisation of the two tested environments. Several key findings emerge from this study.

We identified seven conserved genes important in *S*. Typhimurium establishment on both alfalfa sprouts and glass, highlighting these shared genes as critical for *S*. Typhimurium colonisation of diverse surfaces. These genes belong to various functional categories, including NADH hydrogenase synthesis (*nuoA* and *nuoB*), fimbrial regulation and production (*fimA* and *fimZ*), LPS O-antigen synthesis (*rfbJ*), iron acquisition (*ybaN*), and stress responses (*rpoS*). Ethanolamine utilisation genes, *eutT* and *eutQ*, were also identified to play an important role in *S*. Typhimurium establishment on both environments, with their disruption aiding colonisation of both surfaces. Notably, ethanolamine signalling has been reported to aid *S*. Typhimurium infection of mammalian cells (Srikumar and Fuchs, 2011). The identification of these conserved genes underscores their significance in surface colonisation, regardless of the surface material and shows a core requirement for different pathways providing diverse functions in colonisation to surfaces.

There were also significant differences between the two conditions. Flagella biosynthesis was found to affect colonisation of biotic and abiotic surfaces differently in our study. We showed that aflagellated mutants (*ΔflhDC* and *ΔflhA*) exhibit significantly enhanced glass colonisation at the early stages of colonisation (24 hours) but perform significantly worse on alfalfa. However, with time, these mutants regain their ability to grow on alfalfa. This demonstrates the potential role of the flagellum for initial stages of adhesion to the roots. We know that flagellar motility is essential for initial host colonisation in several bacterial species (Haefele and Lindow, 1987, Van de Broek et al., 1998). This contrasts with the prediction from the TraDIS-*Xpress* results which identified the system as under selective pressure but suggested mutants would have a benefit. This highlights the complexity of the role of flagella at different stages of colonisation but also is likely to reflect the differences between comparing fitness of mutants within a pool to as a monoculture where functions cannot be provided by neighbouring cells (Rossez et al., 2015).

We also found pathways involved in *S*. Typhimurium establishment on alfalfa seedlings that were not involved in biofilm formation on glass. Notably, genes related to type III secretion regulation (*sirC*), Fe-S cluster assembly (*iscA*), curcumin degradation (*curA*), and copper tolerance (*cueR*) confer a competitive advantage to *S*. Typhimurium during colonisation of alfalfa. Deletion of *cueR* reduced the ability of *S*. Typhimurium to colonise plants but had no effect on glass, demonstrating a conditional importance between surfaces. Metals play an important role in plant-pathogen interactions (Fones and Preston, 2013) with copper being commonly deployed as a host antimicrobial. Regulating the expression of copper export through *cueR* is clearly beneficial for colonisation and establishment on a plant. Deletion of *iscA* reduced colonisation of both glass and plant surfaces, and there was no difference in colonisations seen in Δ*sirC* or Δ*curA* deletion mutants. TraDIS-*Xpress* can determine very small changes in competitive fitness that may not always be seen in culture-based assays, therefore further characterisation is needed to determine how these genes affect plant colonisation.

In conclusion, this research provides a comprehensive understanding of the genetic determinants that influence *S*. Typhimurium colonisation and establishment on diverse surfaces. The findings emphasise the role of specific genes mediating adhesion, metal tolerance and stress responses in different stages of *S*. Typhimurium colonisation of fresh produce. This knowledge advances our understanding of *Salmonella* pathogenesis and host-microbe interactions and may have implications for controlling *Salmonella* colonisation and infection.

## Materials and Methods

### Alfalfa seed sterilisation and germination

Alfalfa seeds were sterilised by immersion in a Falcon tube containing 20 mL of 70% ethanol for 30 seconds, followed by three sequential rinses with 20 mL sterile water. Subsequently, the seeds were treated with 5% sodium hypochlorite (20 mL) for 3 minutes on a rolling platform. Three subsequent washes in water were carried out. For germination, sterilised seeds were transferred to square agar plates (20 mL) containing Murashige-Skoog (MS) agar medium. These seeds were positioned with sufficient spacing to allow for three days of germination, reaching an approximate size of 1 cm. Following germination, the seedlings were transferred to fresh MS plates and infected with *S*. Typhimurium strain. Adequate seedlings were included in the process to enable replication for experimental purposes.

### Visualisation of *Salmonella* on alfalfa seedlings

Three-day-old alfalfa seedlings were infected with *S*. Typhimurium tagged with the *lux* operon (*14028S::lux*). A total of 20 μL was evenly distributed in two separate applications along the roots, ensuring even distribution, with the bacterial density normalized to an optical density (OD600nm) of 0.2. The seedlings were incubated at 20°C throughout the experiment’s duration. Visual assessments were conducted at multiple time points following infection, specifically at 3 hours, 24 hours, 48 hours, and 72 hours post-infection. The Biorad Gel Documentation system (ChemiDoc™ MP) was used to capture chemiluminescence, and in addition to this, a colorimetric image was captured and superimposed over the luminescent image for a comprehensive analysis of *S*. Typhimurium presence on the seedlings. Images were analysed using ImageJ by generating merged channel images. Increase in pixel intensity is indicated by change in colour from blue to red, indicating higher cell density of the *14028S::lux* strain.

### Competition assays on glass and on alfalfa seedlings

For competition in alfalfa seedlings, three-day-old seedlings were infected with 10 μL of *S*. Typhimurium tagged with lacZ (*14028S::lacZ*) in a 1:1 ratio with deletion mutants, all adjusted to a final OD of 0.02 in 10mM MgCl2. Infected seedlings were subsequently transferred to fresh MS plates and incubated at 20°C. After 24, 48, and 72 hours post-infection, three seedlings per timepoint were homogenised using a plastic pestle in PBS and then serially diluted in PBS. The dilutions were spotted on LB-agar plates supplemented with 40 μg/ml X-gal (-Bromo-4-chloro-3-indolyl β-D-galactopyranoside) and 1mM IPTG (Isopropyl β-D-1-thiogalactopyranoside).

For competition on glass beads, glass beads suspended in 5 mL of LB-NaCl were inoculated with 50 μL of selected strains mixed with *14028S::lacZ* in a 1:1 ratio, normalized to a final OD of 0.02. After incubation, three beads were recovered at 24, 48, and 72 hours, washed in PBS to eliminate planktonic growth, and the biofilm cells were recovered by vortexing in PBS. The recovered cells were serially diluted and spotted on LB agar plates supplemented with 40 μg/ml X-gal and 1mM IPTG.

### *Salmonella* cell recovery from alfalfa

To perform cell counting, a *S*. Typhimurium strain tagged with *lac*Z (*14028S::lacZ*) (Holden et al., 2020) was used. Seedlings were infected with this strain following the previously described procedure. Cell isolation was carried out at 6 hours, 24 hours, 48 hours, and 72 hours post-infection. For each time point, three seedlings were processed. Individual seedlings were homogenised in 500 μL of sterile PBS using a plastic pestle. The resulting samples underwent serial dilution, and these dilutions were plated on lysogeny broth (LB) agar plates. These plates were supplemented with 40 μg/mL X-Gal and 1 mM IPTG to allow blue-white screening of colonies. The prepared plates were incubated at 37°C overnight. Following overnight incubation, colony-forming units (CFU) were counted to give a measure of bacterial load per seedling. Each time point included at least three technical replicates and three biological samples, ensuring robust and reliable quantification of *S*. Typhimurium populations.

### TraDIS-*Xpress* library preparation, sequencing and data analysis

Three-day-old alfalfa seedlings, grown on MS agar, were infected at the shoot-root junction with a 5 μL droplet of a *S*. Typhimurium transposon mutant library (described by Holden et al. (2022), normalised to an OD_600nm_ of 0.01 with 1 mM IPTG to induce transcription from the transposon-located promoter. Seedlings were then allowed to grow at 30 °C before sampling following 24, 48 and 72 hours growth. Ten seedlings were processed per timepoint and were homogenised in 1 mL of sterile PBS using a plastic pestle. Samples were filtered through 5 μm syringe filters to isolate bacterial cells and eliminate plant cell contamination. Genomic DNA was extracted from these cells following the protocol described by Trampari et al. (2021) Genomic DNA was extracted from these cells following the protocol described by Trampari et al. (2021). A Museek DNA fragment library preparation kit (ThermoFisher) was used to tagment genomic DNA and was then purified with AMPure XP beads (Beckman Coulter). DNA fragments were amplified using customised primers that anneal to the tagmented ends and biotinylated primers that anneal to the transposon. These PCR products were purified as previously and biotinylated DNA was incubated for 4 hours with streptavidin beads (Dynabeads® kilobaseBINDER™, Invitrogen) to capture only DNA fragments containing the transposon. These fragments were amplified using barcoded sequencing primers that anneal to the tagmented ends and to the transposon (Yasir et al., 2020). DNA fragments were then purified and size-selected using AMPure beads. Fragment length was quantified using a Tapestation (Aligent) and sequenced on a NextSeq500 using the NextSeq 500/550 High Output Kit v2.5 with 75 cycles.

Fastq files were aligned to the *S*. Typhimurium 14028*S* reference genome (CP001363, modified to include chromosomally integrated *lacIZ*) using BioTraDIS (version 1.4.3) (Barquist et al., 2016). Significant differences (*p* < 0.05, after correction for false discovery) in insertion frequencies between planktonic and biofilm conditions at each time point were found using BioTraDIS and AlbaTraDIS (version 1.0.1) (Page et al., 2019).

### Mutant creation and phenotypic validation

The luminescent S. Typhimurium strain tagged with lux was created following the gene doctoring protocol (Lee et al., 2009) using vectors created by Holden et al. (2020). Single gene deletion mutants were made following the gene doctoring protocol. The luminescent S. Typhimurium strain tagged with lux was created following the gene doctoring protocol (Lee et al., 2009) using vectors created by Holden et al. (2020). Single gene deletion mutants were made following the gene doctoring protocol (Lee et al., 2009) using plasmids constructed via Golden Gate assembly (Thomson et al., 2020). Mutants were validated by whole genome sequencing to confirm loss of the gene of interest. Primers for mutant construction are listed in supplementary table 2.

