## Supplementary materials for "Investigating how *Salmonella* colonise alfalfa using a whole genome screen"

**Supplementary data**

**Table S1:** Genes determined by TraDIS-*Xpress* to be important for biofilm formation by *Salmonella enterica* serovar Typhimurium on alfalfa sprouts over time. Log-fold change between biofilm and planktonic conditions at each time point are only shown for genes where there are differences in insertion frequency inside the coding region. Where the plot files generated by BioTraDIS show a difference in insertion frequency upstream or downstream of a gene, log-fold change cannot easily by quantified and therefore the effect has been described in the column titled ‘observed change’. Significant differences in insertion frequencies have been manually verified with the plot files generated by BioTraDIS.

<table>
<tr><th>Pathway</th><th>Gene</th><th>Time point</th><th>Log-fold change</th><th>Observed change from planktonic control</th></tr>
<tr><td rowspan="8">LPS</td><td>rfaE</td><td>24h</td><td>-10.82</td><td>Fewer insertions</td></tr>
<tr><td>rfbC</td><td>72h</td><td>-1.05</td><td>Fewer insertions</td></tr>
<tr><td>rfbF</td><td>72h</td><td>-2.90</td><td>Fewer insertions</td></tr>
<tr><td>rfbH</td><td>48h<br>72h</td><td>-2.18<br>-2.68</td><td>Fewer insertions</td></tr>
<tr><td>rfbI</td><td>72h</td><td>-1.34</td><td>Fewer insertions</td></tr>
<tr><td>rfbJ</td><td>48h</td><td>-10.28</td><td>Fewer insertions</td></tr>
<tr><td>rfbV</td><td>48h<br>72h</td><td>-12.13<br>-10.49</td><td>Fewer insertions</td></tr>
<tr><td>galU</td><td>24h</td><td>-11.24</td><td>Fewer insertions</td></tr>
<tr><td rowspan="6">Respiration</td><td>nuoA</td><td>24h<br>72h</td><td>-10.07<br>-9.64</td><td>Fewer insertions</td></tr>
<tr><td>nuoB</td><td>24h</td><td>-9.70</td><td>Fewer insertions</td></tr>
<tr><td>eutT</td><td>24h<br>48h<br>72h</td><td>2.99<br>3.52<br>2.69</td><td>More insertions</td></tr>
<tr><td>pykF</td><td>24h</td><td>-3.50</td><td>Fewer insertions</td></tr>
<tr><td>aceE</td><td>48h</td><td>-11.00</td><td>Fewer insertions</td></tr>
<tr><td>gntR</td><td>72h</td><td>-2.40</td><td>Fewer insertions</td></tr>
<tr><td rowspan="6">DNA housekeeping</td><td>xseA</td><td>72h</td><td>3.17</td><td>More insertions</td></tr>
<tr><td>yoaA</td><td>72h</td><td>-2.62</td><td>Fewer insertions</td></tr>
<tr><td>rnhA</td><td>48h</td><td>-10.02</td><td>Fewer insertions</td></tr>
<tr><td>ung</td><td>48h</td><td>-10.52</td><td>Fewer insertions</td></tr>
<tr><td>STM14_1174</td><td>24h</td><td>-9.16</td><td>Fewer insertions</td></tr>
<tr><td>STM14_4641</td><td>24h<br>48h<br>72h</td><td>1.84<br>3.26<br>3.97</td><td>More insertions</td></tr>
<tr><td rowspan="3">Stress response transcription factors & regulators</td><td>rpoS</td><td>72h</td><td>-3.42</td><td>Fewer insertions</td></tr>
<tr><td>rpoE</td><td>24h<br>48h<br>72h</td><td>-9.51<br>-9.71<br>-9.87</td><td>Fewer insertions</td></tr>
<tr><td>crl</td><td>72h</td><td>-1.53</td><td>Fewer insertions</td></tr>
</table>

| | | | | |
|---|---|---|---|---|
| | *yaiB/ iraP* | 48h<br>72h | -1.03<br>-2.31 | Fewer insertions |
| Flagella | *flhA* | 72h | 2.01 | More insertions |
| | *flgG* | 72h | 2.08 | More insertions |
| | *fliE* | 72h | 2.21 | More insertions |
| | *fliS* | 72h | 2.69 | More insertions |
| Fe-S cluster assembly | *iscA* | 24h | -10.18 | Fewer insertions |
| | *nifS* | 72h | -9.45 | Fewer insertions |
| | *yhgI/ nfuA* | 24h | -10.63 | Fewer insertions |
| Copper tolerance | *cueR* | 24h<br>72h | -9.22<br>-10.14 | Fewer insertions |
| | *scsD* | 72h | -9.64 | Fewer insertions |
| Protein transport and folding | *surA* | 24h<br>48h<br>72h | -7.85<br>-11.08<br>-11.23 | Fewer insertions |
| | *smpA/ bamE* | 48h | -10.46 | Fewer insertions |
| | *yacA/ secM* | 48h<br>72h | -9.55<br>-9.82 | Fewer insertions |
| Fimbriae | *fimA* | 48h<br>72h | | Increased expression relative to planktonic control |
| | *fimZ* | 24h | -1.04 | Fewer insertions |
| Cell envelope | *nlpD* | 72h | -3.08 | Fewer insertions |
| | *pgpA* | 72h | -10.01 | Fewer insertions |
| Type III secretion system | *sirC* | 24h | -9.20 | Fewer insertions |
| Curcumin degradation | *yncB/ curA* | 72h | -0.94 | Fewer insertions |
| Thiamine biosynthesis | *thiL* | 72h | -9.58 | Fewer insertions |
| MFS phosphate transporter | *glpT* | 24h | -3.17 | Fewer insertions |
| Protease specificity-enhancing factor | *sspB* | 24h | -6.86 | Fewer insertions |
| Iron storage | *bfd* | 48h | -9.96 | Fewer insertions |
| Biofilm protein | *yjgK/ tabA* | 72h | -2.18 | Fewer insertions |
| Transcription | *slpA/fkpB* | 48h | -10.04 | Fewer insertions |
| Lipid hydrolase | *ychK/rssA* | 72h | -1.89 | Fewer insertions |
| Oxidoreductase | *STM14_2022* | 48h | -10.41 | Fewer insertions |
| Unknown | *yaaY* | 24h | -10.18 | Fewer insertions |
| | *ybaN* | 72h | -9.51 | Fewer insertions |
| | *ybjC* | 48h<br>72h | -9.55<br>-9.99 | Fewer insertions |
| | *ydiZ* | 24h | -9.12 | Fewer insertions |

| | | | | |
|---|---|---|---|---|
| | *yqjF* | 24h | -10.49 | Fewer insertions |
| | *STM14_0076* | 24h | -9.38 | Fewer insertions |
| | *STM14_0143* | 72h | -9.49 | Fewer insertions |
| | *STM14_0487* | 24h | -9.32 | Fewer insertions |
| | *STM14_0526* | 72h | -11.26 | Fewer insertions |
| | *STM14_0643* | 24h<br>72h | 2.84<br>0.81 | More insertions |
| | *STM14_1102* | 72h | -9.54 | Fewer insertions |
| | *STM14_1108* | 72h | -9.65 | Fewer insertions |
| | *STM14_1993* | 24h | -9.35 | Fewer insertions |
| | *STM14_2059/ yciZ* | 72h | -11.22 | Fewer insertions |
| | *STM14_2117* | 24h | -9.19 | Fewer insertions |
| | *STM14_3249* | 24h | -10.15 | Fewer insertions |
| | *STM14_4582* | 24h | -9.74 | Fewer insertions |
| | *STM14_5469* | 24h<br>48h<br>72h | -9.90<br>-9.86<br>-10.20 | Fewer insertions |
| | *STM14_5479* | 72h | -9.97 | Fewer insertions |

**Figure S1:** Insertion frequency per gene in an *S*. Typhimurium transposon mutant library colonising alfalfa plants (x-axis) compared to planktonic conditions (y-axis) after 1 day (24 hours), 2 days (48 hours) and 3 days (72 hours). Black points show the insertion frequency per gene for each replicate to display the variation between replicates, and coloured points show the mean insertion frequency per gene of the biofilm condition compared to the planktonic condition.

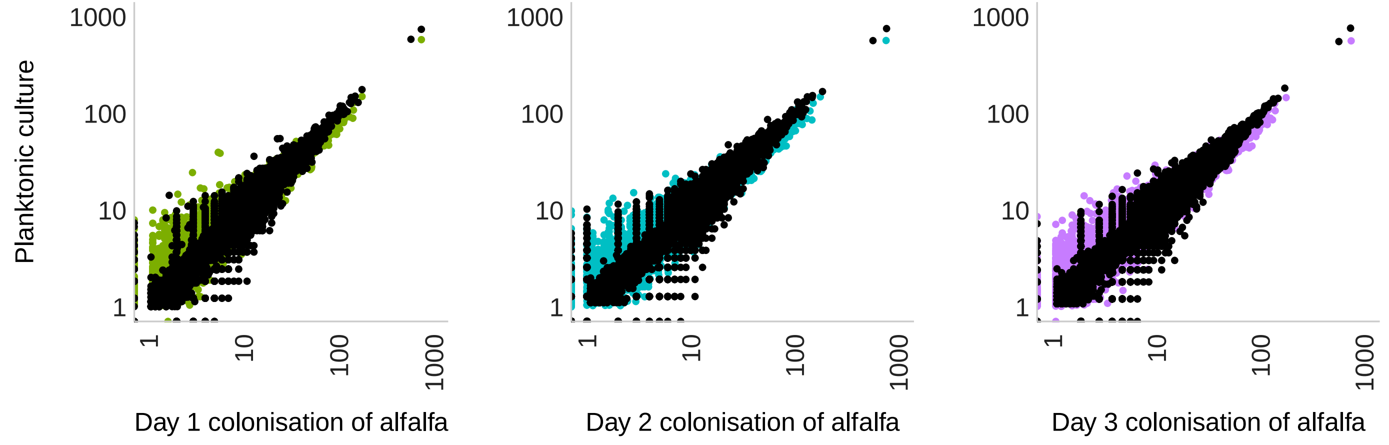


**Table S2:** Primers used in this work. Customised indices for TraDIS-*Xpress* sequencing library preparation can be found in the supplementary material of Yasir et al (2020; doi: 10.1101/gr.254391.119)

| **Name** | **Sequence** | **Function** |
|---|---|---|
| pdoc-K-glms-lux-For | AATGCGCTCGAGGTTGCAAATTTTTCAACATTTTATACACTACGAAAACCATCGCGAAAGCGAGTTTTGGATTTAAGAAGGAGATATACATATGACTAAAAAAATTTCATTCATTATTAA | Amplification of lux operon from pUC18-mini-Tn7-lux under the control of the *acpP* promoter |
| pDOC-K-glms-lux-Rev | GCTCAGAAGCTTTCAACTATCAAACGCTTCG | |
| flhA HR1 fwd | GGGGGTCTCGCTACGCGTTTCTATTTATTCAAAAAGAGAGTCAGGTCT | Amplification of homologous regions upstream and downstream of *flhA* for its deletion in *S.* Typhimurium |
| flhA HR1 rev | GGGGGTCTCTCTCCTTTCCGATAACCGTCATATCCGCA | |
| flhA HR2 fwd | GGGGGTCTCTCGCTGTCGATTTCAGGTTGCTGGGC | |
| flhA HR2 rev | GGGGGTCTCATCGTAAGAGAGCGAAGGCGATCCG | |
| flhDC HR1 fwd | GGGGGTCTCGCTACATTTCACTCTCTTTTGGATTTTCAATATCGCG | Amplification of homologous regions upstream and downstream of *flhDC* for its deletion in *S.* Typhimurium |
| flhDC HR1 rev | GGGGGTCTCTCTCCCCGATATTATTCCACAACTGCTGGATGAA | |
| flhDC HR2 fwd | GGGGGTCTCTCGCTTTGATGTCATAAATGTGTTTTAGCAACTCGG | |
| flhDC HR2 rev | GGGGGTCTCATCGTCTGTTATCTATTATCCTGGCGTTATTTTAACAGAGAG | |
| cueR HR1 fwd | GGGGGTCTCGCTACGTGACACAACTGGCGCAGC | Amplification of homologous regions upstream and downstream of *cueR* for its deletion in *S.* Typhimurium |
| cueR HR1 rev | GGGGGTCTCTCTCCATGGCTTTGCTGGTTAAACCGGT | |
| cueR HR2 fwd | GGGGGTCTCTCGCTTTGATAATCTTTCCGGCTGCTGTCA | |
| cueR HR2 rev | GGGGGTCTCATCGTATAGCGATGATTCTGGTGCATCCG | |
| sirC HR1 fwd | GGGGGTCTCGCTACTTCATCAAGCGTTTCACCGTTGAAC | |

| | | |
|---|---|---|
| sirC HR1 rev | GGGGGTCTCTCTCCTGTACATCATCATTAAACTCGCCACCA | Amplification of homologous regions upstream and downstream of *sirC* for its deletion in *S.* Typhimurium |
| sirC HR2 fwd | GGGGGTCTCTCGCTGAGAGCGCAACACAGATAAGATGAAGC | |
| sirC HR2 rev | GGGGGTCTCATCGTGGCCTGCGAGCGACC | |
| iscA HR1 fwd | GGGGGTCTCGCTACCGTATGTTGCTCGCTGGCCAG | Amplification of homologous regions upstream and downstream of *iscA* for its deletion in *S.* Typhimurium |
| iscA HR1 rev | GGGGGTCTCTCTCCACCCGAATGTGAAAGATGAGTGTGGTT | |
| iscA HR2 fwd | GGGGGTCTCTCGCTAGGAAGGTATTAACTCGCGCTGC | |
| iscA HR2 rev | GGGGGTCTCATCGTATCATTAATCGGTATCGGAATCAGGAGAAT | |
| curA HR1 fwd | GGGGGTCTCGCTACATTTTTAGTGATAAGCCTTGCGCCT | Amplification of homologous regions upstream and downstream of *curA* for its deletion in *S.* Typhimurium |
| curA HR1 rev | GGGGGTCTCTCTCCCGGGGAAGAACTTTGGCAAAGT | |
| curA HR2 fwd | GGGGGTCTCTCGCTTTGGTCTGTTGCTTCATTCTATTCTCCT | |
| curA HR2 rev | GGGGGTCTCATCGTTGCGCCGGATGAACATAAACC | |
| rfbJ HR1 fwd | GGGGGTCTCGCTACAGACATGAGGTAAAAAAGAGCATTCTGG | Amplification of homologous regions upstream and downstream of *rfbJ* for its deletion in *S.* Typhimurium |
| rfbJ HR1 rev | GGGGGTCTCGCTCCGGTTATGAGATTTTCATGATCTTTTAATAAATAAATCGTTAACAAA | |
| rfbJ HR2 fwd | GGGGGTCTCTCGCTGGTCATCGCAATCACCAGATAGAATAAATTG | |
| rfbJ HR2 rev | GGGGGTCTCATCGTCCCACCTCGGATATAATCTCAAAATCACG | |
